# Hydrogel-Embedded Precision-Cut Lung Slices Recapitulate Fibrotic Gene Expression and Enable Therapeutic Response Evaluation

**DOI:** 10.64898/2026.03.24.714004

**Authors:** Alicia E. Tanneberger, Rachel Blomberg, Tvishi Yendamuri, Haley Noelle, Jeffrey G. Jacot, Janette K. Burgess, Chelsea M. Magin

## Abstract

Precision-cut lung slices (PCLS) retain the native cells and extracellular matrix that contribute to the structural and functional integrity of lung tissue. This technique enables the study of cell-matrix interactions and is particularly useful for pre-clinical pharmacological studies. More specifically, PCLS are widely used to model the complex pathophysiology of pulmonary fibrosis, an uncurable and progressive interstitial lung disease. Current *ex vivo* pulmonary fibrosis models expose PCLS to pro-fibrotic biochemical cues over a short timeframe (hours to days) and quickly collect samples for analysis due to viability concerns. This condensed timeline is a limitation to understanding chronic disease mechanisms. To extend the utility of *ex vivo* pulmonary fibrosis models, PCLS were embedded in engineered hydrogels and exposed to pro-fibrotic biochemical and biophysical cues. Hydrogel-embedded PCLS maintained greater than 80% total cell viability over 3 weeks in culture. Gene expression patterns in samples exposed to pro-fibrotic cues matched trends measured in human fibrotic lung tissue. Finally, treatment with Nintedanib, a Food and Drug Administration approved pulmonary fibrosis drug, moderately reduced fibroblast activation and influenced epithelial cell differentiation. Collectively, these results show that hydrogel-embedded PCLS models of pulmonary fibrosis extend our ability to study fibrotic processes *ex vivo* and, when applied to human tissues, present a new approach methodology for studying lung disease and treatment.

## Introduction

Idiopathic pulmonary fibrosis (IPF), one of the most frequent forms of interstitial lung disease, has a current median survival rate of only 2-3 years after diagnosis (1). Until very recently, only two Food and Drug Administration (FDA)-approved drugs (Pirfenidone and Nintedanib) existed to manage IPF symptoms, but neither showed substantial improvement in patient survival rates (2, 3). New drug approval is expensive, time consuming, and must pass multiple rigorous development stages. Approximately 90% of successful preclinical drugs fail to show comparable clinical efficacy in human pulmonary fibrosis (4). As such, the only cure for many end-stage patients is lung transplantation, yet the number of individuals needing an organ transplant far outweighs the number of available organs (5). Therefore, there remains a critical need for improved *ex vivo* lung tissue models that systematically investigate dynamic biological fibrotic processes and advance the drug development timeline.

Precision-cut lung slices (PCLS) are a versatile *ex vivo* model that preserves the native three-dimensional (3D) tissue architecture, cellular heterogeneity, and extracellular matrix (ECM) of lung tissues. Historically, PCLS were primarily used to study airway contractions, but this tool has since gained prominence in distal lung tissue research (6, 7). By maintaining structural and cellular integrity, PCLS provide a physiologically relevant platform for studying lung injury, repair, fibrosis, and regenerative therapies, spanning fundamental biological research to high-throughput drug screening. Overwhelming new evidence shows that culturing cells in 3D most closely mimics *in vivo* conditions, helps retain *in vivo* cellular physiology and molecular characteristics, and improves translational efforts (8–12). Importantly, the thin tissue slice allows for real-time imaging of dynamic processes, such as epithelial migration, immune cell behavior, and tissue remodeling (13, 14). Live imaging in PCLS has revealed the dynamic behavior of epithelial cells during alveologenesis and highlighted the role of innate immunity in lung repair following viral infections (15).

PCLS are commonly exposed to pro-fibrotic stimuli, such as transforming growth factor beta (TGF-β) to model fibrosis progression (16–18). The incorporation of a fibrosis cocktail (FC), typically composed of TGF-β in combination with platelet-derived growth factor, tumor necrosis factor alpha, and lysophosphatidic acid, provides a strong exogenous stimulus that promotes robust fibrotic activation and has been broadly implemented across *ex vivo* models (16, 17, 19, 20). These efforts have facilitated the identification of critical pathways involved in ECM remodeling and repair, providing critical insights into potential therapeutic targets (21). Furthermore, transcriptomic analyses of PCLS have uncovered cellular and molecular responses during early fibrotic changes, offering a platform for investigating the progression of lung fibrosis in a controlled environment (18, 22, 23). Recent studies also highlighted the value of PCLS in high-throughput drug screening platforms, where these models were used to assess the efficacy and safety of anti-fibrotic therapies and other interventions (15, 23).

Despite the advanced and highly translational nature of PCLS, traditional culture systems remain limited by short-term viability. Culture lifespans of 7-10 days substantially restrict the ability to investigate long-term processes such as lung injury and repair (24–26). To address these limitations we introduced hydrogel-embedded PCLS, a new approach methodology, that encapsulates PCLS within poly(ethylene glycol) norbornene (PEGNB)-based hydrogel biomaterials to extend *ex vivo* viability for up to six weeks (26, 27). PEGNB hydrogels are a highly defined and reproducible alternative to many commercial ECM products, which often exhibit substantial batch-to-batch variability (e.g., Matrigel). As a synthetic biomaterial, PEGNB hydrogels offer a versatile 3D culture platform with tunable composition and mechanical properties, advantages that are difficult to achieve with naturally derived matrices (28). In particular, PEGNB hydrogels can be precisely engineered to recapitulate the mechanical stiffness of both healthy and diseased lung tissue (19, 27, 29). Healthy lung tissue exhibits an elastic modulus (E) in the range of 1-5 kPa, while fibrotic lung tissue is characterized by significantly increased stiffness, frequently exceeding 10 kPa (28, 30).

PEGNB is a phototunable hydrogel backbone that can be combined with a wide range of crosslinkers including molecules that are degradable by matrix metalloproteinase (MMP) enzymes secreted by the cells contained within the constructs. Cell-adhesive peptide-mimics, including sequences representing fibronectin (CGRGDS), collagen (CGFOGER), or laminin (CGYIGSR), can also be incorporated into the hydrogel formulation to enhance cell binding (19, 27, 31–34). Similarly, targeted MMP-degradable sequences, relevant to lung tissue such as MMP9, can be integrated into the hydrogels to support cell remodeling (19, 29, 32, 35). By leveraging click chemistry and dual-stage polymerization systems, biomaterial properties can be spatially and temporally controlled by user input (28, 33, 36–39). For instance, soft hydrogels containing tyrosine residues can be dynamically stiffened using ruthenium crosslinking chemistry, which mimics the increased ditryosine crosslinking observed in fibrotic lungs, a key feature of pathophysiological ECM remodeling (14, 40).

Prior work by our group highlights how hydrogel-embedded PCLS are a powerful platform that enables mechanistic and longitudinal studies difficult to perform in whole-animal models (27, 41). Compared with *in vivo* experiments, PCLS reduce the number of animals required per study and expand experimental throughput by allowing multiple conditions to be tested from a single lung (7, 16, 42, 43). Although hydrogel-embedded human PCLS are the ideal final tool for investigating lung disease and treatment, human studies are limited by inter-donor and tissue availability. Mouse PCLS have historically served as a suitable alternative for many experiments and are well-studied for proof-of-concept studies (25, 27, 44, 45). The bleomycin mouse model of pulmonary fibrosis, for example, is widely established as a well-recognized pulmonary fibrosis model (46–50). Here, control murine PCLS were embedded in PEGNB hydrogels, then systematically exposed to pro-fibrotic biochemical cues and dynamic mechanical stiffening to dissect how biophysical and biochemical signals cooperatively drive fibrotic remodeling over extended culture durations.

## Materials and Methods

### Animal Use

All animal use was carried out as approved by the Institutional Animal Care and Use Committee at the University of Colorado Anschutz (protocol 1002). Each experimental condition used equal numbers of PCLS generated from male and from female mice to acknowledge sex as a biological variable.

### PCLS Procurement and Processing

Adult (8-15 weeks) C57BL/6J mice were sacrificed by a lethal dose of ketamine and xylazine, delivered by intraperitoneal injection, and secondly by exsanguination. To clear the lungs, a cardiac perfusion of 10 mL of sterile phosphate buffered saline (PBS) was injected into the right ventricle of the heart. Lungs were inflated with 1.5 weight percentage (wt%) low melting point agarose dissolved in sterile PBS through the trachea and covered with ice for 10 min to solidify the agarose. The lungs were then manually extracted and placed in PCLS medium (0.1% Penicillin/Streptomycin/Fungizone/Gentamycin (P/S/F/G) antibiotics/antimycotics and 0.1% charcoal stripped fetal bovine serum (CS-FBS) in Dulbecco’s Modified Eagle Medium (DMEM)/Nutrient Mixture F-12).

Individual lobes were sliced into 500-μm whole tissue slices using a vibratome (Campden Instruments) at a speed of 12 mm s^-1^. Standardized punches were created using a 4-mm biopsy punch (Fisher Scientific). Tissue punches were initially incubated in PCLS medium at 37°C and 5% CO₂ for 30 minutes. Following this, the medium was refreshed and the incubation step was repeated twice, resulting in three consecutive 30-minute washes. Punches were then cultured overnight in 24-well plates within PCLS medium (37°C and 5% CO_2_).

### PEGNB Synthesis

PEGNB was synthesized as previously described from an eight-arm, 10 kg/mol PEG-hydroxyl macromer (PEG-OH) (19, 29, 32, 35). In brief, the lyophilized PEG-OH (5 g, JenKem Technology) was dissolved in anhydrous dichloromethane (DCM; 35 mL, Sigma-Aldrich) and reacted with both 4-Dimethylaminopyridine (DMAP; 0.24 g, 0.002 mol, Acros Organics) and pyridine (1.61 mL, 0.02 mol, Sigma Aldrich). In a second reaction flask, norbornene-2-carboxylic acid (4.9 mL, 0.04 mol, Acros Organics) was added dropwise to N,N’-Dicyclohexylcarbodiimide (DCC; 4.13 g, 0.02 mol, Fisher Scientific) dissolved in anhydrous DCM. After 30 min of stirring, the second reaction mixture was filtered through Celite 545 (EMD Millipore, cat. #CX0574-1), added to the first flask, and left to react for 48 h. Undesired byproducts were removed through a series of washes. The organic product was precipitated out with cold diethyl ether (Fisher Scientific) and then concentrated with a rotary evaporator. After removing the diethyl ether, the precipitate was vacuum dried and dialyzed over 72 h. After dialysis, the product was collected and lyophilized (∼ 0.1 mBar, ∼ -80°C) to obtain a solid white powder.

End-group functionalization and purity were confirmed with nuclear magnetic resonance (NMR) spectroscopy. A Bruker DPX-400 FT NMR spectrometer collected the ^1^H NMR spectrum of the product using 184 scans and a 2.5 s relaxation time. Synthesis products with 89% or greater functionalization were used throughout these experiments, and chemical shifts for protons (^1^H) were recorded relative to deuterochloroform as parts per million (ppm) (**Supplemental Figure 1**).

### Hydrogel Preparation and PCLS Embedding

A hydrogel precursor solution was prepared by combining 3.75-4 wt% PEGNB with a matrix metalloproteinase-9 (MMP9)-degradable crosslinker (Ac-GCRD-VPLSLYSG-DRCG-NH2, 9.6-10.2 mM, GL Biochem) in a 0.7 ratio of thiols-to-norbornenes and the cellular adhesion peptides CGRGDS (0.1 mM; fibronectin mimic, GL Biochem), CGYIGSR (0.2 mM; laminin mimic, GL Biochem), and CGFOGER (0.1 mM; collagen mimic, GL Biochem). The PEGNB, MMP9-degradable crosslinker, CGRGDS, CGYIGSR, and CGFOGER were all dissolved in PCLS medium to create stock solutions. Then, components were mixed to create a final hydrogel precursor solution with the concentrations listed above. To confirm that the pH of the hydrogel precursor solution was neutral (between 7 and 8) prior to any cell exposure, 2 μL of the hydrogel precursor solution was pipetted onto pH indictor strips. One MMP-9 degradable peptide batch was acidic when received from the supplier. To neutralize the overall hydrogel precursor solution pH prior to embedding, the MMP9-degradable peptide was resuspended in a solution of 10% v/v 1 M sodium hydroxide and 90% v/v PCLS medium.

A bottom layer of hydrogel solution (25 μl) was polymerized within a round, 8-mm diameter silicone mold using a light-mediated thiol-ene reaction and the photoinitiator lithium phenyl-2,4,6-trimethylbenzoylphosphinate (2.2 mM, LAP; Sigma Aldrich) (26, 27). Five minutes of ultraviolet (UV) light exposure (365 nm, 10 mW cm^−2^, Omnicure, Lumen Dynamics) facilitated crosslinking. Then, a single PCLS punch was carefully placed onto this base layer, covered with an additional 25 μl of hydrogel precursor solution, and exposed to 5 min of 365 nm, 10 mW cm^−2^ light. The polymerized hydrogel-embedded PCLS constructs were gently transferred into 24-well plates and incubated in PCLS medium at 37°C and 5% CO_2_.

### Pro-Fibrotic Biochemical Cue Exposure

Hydrogel-embedded PCLS were maintained in PCLS medium (37°C, 5% CO_2_) that was changed every 48 h. On days 2, 6, 10, 14, and 18, wells were replenished with 0.5 mL of PCLS medium. Starting on day 4, samples were exposed to a fibrosis cocktail (FC) or vehicle control (VC). The fibrosis cocktail contained 10 ng/ml platelet-derived growth factor AB (PDGF-AB; Thermo Scientific, cat. #PHG0134), 5 ng/ml recombinant transforming growth factor beta (TGF-β; PeproTech, cat. #100-21), and 5 μM 1-Oleoyl Lysophosphatidic Acid (LPA; Cayman Chemical Company, cat. #62215) (16, 20, 51, 52). FC exposure occurred every four days and each well was replenished with medium (0.5 mL/well) containing either the FC or VC (PBS supplemented with 0.1% bovine serum albumin (BSA)) on days 4, 8, 12, 16, 20.

### Hydrogel Stiffening

A Ruthenium Photoinitiator Kit (Sigma Aldrich) enabled the secondary stiffening response in a proportion of the initially soft hydrogels. Stock solutions of ruthenium (0.047 M) and sodium persulfate (SPS, 0.5 M) were prepared with sterile PBS. 10 μL of each component was supplemented into each 1 mL of PCLS medium so the final concentrations of ruthenium and SPS were 0.47 mM and 5 mM SPS, respectively. The samples to be stiffened were incubated with 0.5 mL/well of the prepared medium over 6 h (37°C, 5% CO_2_). After the incubation period, the medium was manually removed and replaced with sterile PBS for approximately 5 min. The constructs to be stiffened were then exposed to 440 nm visible blue light for 5 min. Again, the constructs were rinsed with PBS and then returned to fresh PCLS medium without photoinitiator.

### Rheological Mechanical Characterization

First, hydrogels were prepared for rheological evaluation without PCLS embedding. The hydrogel precursor solution (40 μL) was pipetted between two glass slides covered in parafilm and separated by a 1-mm gasket by polymerized by UV light (365 nm, 10 mW cm^−2^ intensity, Omnicure, Lumen Dynamics) for 5 min. Hydrogels were then swollen in PBS overnight prior to measurement. Hydrogel storage modulus (G’) was measured using an 8-mm parallel plate geometry on a Discovery HR2 rheometer (TA Instruments), following the methods described in references (35, 47).

Briefly, hydrogels were placed onto the Peltier plate set to 37℃ and the geometry was lowered until it contacted the hydrogel surface and registered an axial force of 0.03 N. The G’ plateau was identified by measuring G’ at successive 5% increments of compression until a maximum was reached (29, 35). The G’ plateau for soft hydrogels occurred at 20% compression and at 25% compression for stiffened hydrogels. Subsequently, the hydrogel samples underwent a frequency oscillation with logarithmic sweep of frequencies ranging from 1-100 rad s^-1^ and a strain of 1% under the compression value where G’ plateaued. The elastic modulus (E) was calculated, assuming the hydrogels were incompressible and exhibited bulk-elastic properties with a Poisson’s ratio of 0.5 (26, 28, 34, 53).

### Cryosectioning

Hydrogel-embedded PCLS were pulled from culture and briefly dipped in hematoxylin (1 min) before being transferred into 50% PBS/50% optimal cooling temperature (OCT) compound for 24 h, while hydrated at room temperature in cryomolds. Following, the samples were transferred into 100% OCT for another 24 h, while hydrated at room temperature in cyromolds. The stepwise OCT incubation improved diffusion into the PEGNB hydrogel and helped reduce shattering during cryosectioning. Samples were then flash frozen by submersion in liquid nitrogen-cooled 2-methylbutane and stored at −80°C until cryosectioning. Samples were cryosectioned on a Leica CM1850 cryostat. The frozen blocks were removed from molds and mounted to specimen disks. 10 μm thickness sections were acquired on positively charged glass slides and stored at -20°C.

### Atomic Force Microscopy (AFM)

Glass slides mounted with sections of tissue and hydrogel were thawed in 1x PBS at room temperature (RT) and maintained in PBS throughout analysis. AFM scans were performed on a JPK NanoWizard 4A Atomic Force Microscope mounted on a Zeiss AxioObserver A1 inverted epifluorescent microscope that allowed visualization of the slides and precise positioning of the probe. AFM measurements all used tip C of an MLCT probe (Bruker AFM Probes, Camarillo, CA) calibrated by thermal oscillation.

Force constants were measured at 0.008–0.010 N/m. Force curves were generated in contact force spectroscopy mode using a maximum force of 500 pN, a retract length of 3 μm, and an extend speed of 2.0 μm/s. For each sample, 10 force curves were collected at different locations in each of the tissue and hydrogel regions. The elastic modulus was evaluated for each force curve by fitting the data to a Hertz/Sneddon model with a pyramidal tip radius. Curves with slips, vibrations, or other excessive noise and curves showing a sharp inelastic surface (the underlying glass) were excluded. The experimental data were analyzed in GraphPad Prism software using a two-way ANOVA with Tukey’s multiple comparisons test. Data are presented as relative stiffness using baseline-corrected values for the soft VC day 15 averages.

### Hydrogel-Embedded PCLS Viability

A ReadyProbes Cell Viability Imaging Kit (Thermo Fischer Scientific) quantified total cell viability. Hydrogel-embedded PCLS were incubated in 0.5 mL of live/dead staining medium for 1 h on an orbital shaker (37°C, 5% CO_2_). This staining medium consisted of 1 drop of NucBlue (nuclei) and 1 drop of NucGreen (dead) per mL of PCLS medium. Hydrogel-embedded PCLS exposed to the VC or FC were collected for imaging after 1, 8, and 15 days in culture. Stiffened samples treated with FC ± Nintedanib were also collected on day 22. Following incubation, hydrogel-embedded PCLS were transferred onto a glass slide and covered in PBS to maintain hydration during imaging. A hydrophobic pen confined the PBS to the sample area. Negative controls were prepared identically but omitted the NucGreen stain as lung tissue exhibits significant autofluorescence in the green channel.

Fluorescently stained samples were imaged on an epifluorescent Olympus BX63 upright microscope. 10x z-stack images (100-200 μm) in the DAPI (nuclei) and FITC (dead) channels were acquired at 3-6 distinct locations within each construct. Maximum intensity projections were created and background fluorescence was subtracted using ImageJ software (NIH). Exposure settings were maintained between samples that were directly compared. Threshold limits were set based on negative control images. The StarDist plugin using the versatile (fluorescent nuclei) model measured cell viability by comparing the number of dead cells to the total number of cell nuclei.

### Nascent Extracellular Matrix Deposition Monitoring

For nascent ECM deposition studies, hydrogel-embedded PCLS were transferred to and kept in a slightly modified version of the previously published 25% concentration L-azidohomoalanine (AHA) medium starting on day 2 until day 15, which contained reduced FBS and P/S concentrations (0.1% v/v) compared to the original formulation (**Supplemental Table 1**) (14, 54, 55). Stock solutions were prepared in advanced in accordance with references (14, 54, 55), but the AHA medium was made fresh each day. Sample imaging occurred on days 8 and 15, which corresponded to the day of stiffening and one-week post-stiffening respectively. The incorporated methionine analog AHA was fluorescently labeled with AZDye 488 DBCO (Vector Laboratories, CCT-1278-1) using copper-free cycloaddition click chemistry (14, 54, 55).

In brief, the samples were manually extracted from the embedding hydrogel and then transferred into 300 μL of washing buffer (2% BSA in PBS) for 5 min at RT (54). Next, samples were submerged in 300 μL of DBCO staining solution (30 μM DBCO in 2% BSA in PBS) (54) that contained a 1:1000 dilution of CellTracker Orange CMTMR Dye (Thermo Scientific; C2927) and 1 drop/mL of NucBlue Live ReadyProbes (Thermo Scientific; R37605) for an additional 1 h at 37 ℃ (54). Meanwhile, negative control samples remained in washing buffer that contained only 1 drop/mL of NucBlue Live ReadyProbes. Following this, samples were rinsed with 300 μL of washing buffer three times, with 5 min incubation periods at RT. 4% v/v paraformaldehyde in PBS was then used to fix the samples for 30 min at RT. Again, the samples were rinsed three times with washing buffer using 5 min incubation periods in between, and then finally were stored at 4 ℃ for up to one week while protected from light.

Similarly, fluorescently stained samples were imaged on the epifluorescent Olympus BX63 upright microscope. Z-stack images (100-200 μm) in the DAPI (blue, nuclei), FITC (green, nascent protein), and TRITC (red, cytoplasm) channels were acquired at 10x for 3 distinct locations within each construct. Maximum intensity projections were created and background fluorescence was subtracted using ImageJ software (NIH). Exposure settings were maintained between samples that were directly compared. Threshold limits were set based on negative control images. Nascent ECM protein was quantified as the percentage area coverage of fluorescence within the green channel.

### Assessment of Gene Expression

PCLS tissue was extracted from the hydrogel using tweezers and placed in a microcentrifuge tube with 700 μL of TRIzol Reagent (Fisher Scientific). Between one and three PCLS per experimental condition were pooled together into a single microcentrifuge tube and sonicated (25% of maximum amplitude) for 5 s to produce an experimental replicate. The samples were then stored at -20°C for up to one month before processing. Afterwards, the samples were thawed and messenger RNA (mRNA) was isolated using Qiagen RNeasy Plus Micro Kits (Qiagen).

Reverse transcription-quantitative polymerase chain reaction (RT-qPCR) assessed gene expression for a variety of different markers: alveolar type II epithelial (ATII) (surfactant protein C (*SFTPC),* lysosome-associated membrane protein 3 (*LAMP3)*), alveolar transitional epithelial (ATII-to-ATI) (keratin 8 (*KRT8),* amphiregulin *(AREG)*), alveolar type I epithelial (ATI) (homeobox only protein X (*HOPX),* advanced glycation end product-specific receptor *(AGER)*), fibroblast activation (collagen 1 alpha chain 1 (*COL1A1),* fibronectin 1 *(FN1),* connective tissue growth factor *(CTGF),* collagen triple helix repeat containing 1 (*CTHRC1),* latent TGF-β binding protein 2 (*LTBP2)*), and hypoxia (hypoxia-inducible factor 1 alpha (*HIF1*α*), lactate dehydrogenase A (LDHA)*). All gene expression was normalized to the housekeeping gene of ribosomal protein S18 (*RPS18*). To normalize the data for statistical analyses, Ct values were transformed by applying the natural log (19, 56) and then relative gene expression was calculated using a 2^−Δ𝐶𝑡^ approach. After statistical analyses were performed, values were untransformed and presented in the figures.

### Response to Anti-Fibrotic Treatment

The FDA-approved anti-fibrotic drug Nintedanib (10 μM, Tocris, cat. #7049) was dissolved in dimethyl sulfoxide. The Nintedanib solution was supplemented into the hydrogel-embedded PCLS medium for stiffened and FC-exposed constructs on days 16, 18, and 20. Stiffened samples that received only the FC components served as controls.

### Statistical Analysis

Data sets were initially assessed for normality within GraphPad Prism using Shapiro-Wilks tests and visual inspection of quantile-quantile (QQ) plots. For data sets that failed both quantitative and visual normality tests (57), natural log transformations (19, 56) were applied prior to performing parametric-based statistical analyses or non-parametric analyses. An unpaired Welch’s t-test calculated statistical significance for rheological measurements. For viability results, t-tests or ordinary one-way ANOVA tests determined statistical significance. Nascent ECM image quantification and AFM results were assessed for statistical significance using a two-way ANOVA test. Lastly, for RT-qPCR results, a natural log transformation was applied and two-way ANOVA tests or unpaired Welch’s t-tests evaluated statistical significance. Data are presented as mean ± standard error of the mean (SEM) and statistical significance was determined by p < 0.05.

## Results

### PEGNB hydrogels recapitulated key aspects of the fibrotic tissue microenvironment

The soft PEGNB hydrogel formulation consisted of an eight-arm, 10 kg/mol macromer that was ≥89% functionalized with norbornene end groups (**Supplemental Figure 1**), an MMP9-degradable peptide crosslinker (containing tyrosine residues), a collagen mimetic peptide (CGFOGER), a laminin mimetic peptide (CGYIGSR, containing tyrosine residues), and a fibronectin mimetic peptide (CGRGDS) (**Figure 1A**). Since this hydrogel formulation contained tyrosine residues through the CGYIGSR peptide and the MMP9-degradable crosslinker sequence, a secondary stiffening mechanism via ditryosine dimerization was theoretically feasible. Using ruthenium chemistry, hydrogel stiffness was expected to increase if ruthenium and SPS were swollen into the hydrogels and then exposed to visible blue light (14, 40, 58). To verify stiffening occurred with the hydrogel alone, rheological measurements were performed after employing these methods. As expected, the secondary chemistry that targeted increasing the number of crosslinks present within the materials enabled the soft hydrogels to significantly increase (p < 0.0001) in stiffness after forming ditryosine dimerizations (14, 40, 58) (**Figure 1C**). Acellular soft hydrogels exhibited an elastic modulus of 2.65 ± 0.25 kPa (falling within the range reported for healthy lung tissue (1–5 kPa) (28, 30)), whereas acellular stiffened hydrogels reached an elastic modulus of 11.81 ± 0.46 kPa (**Figure 1C**).

**Figure 1.**
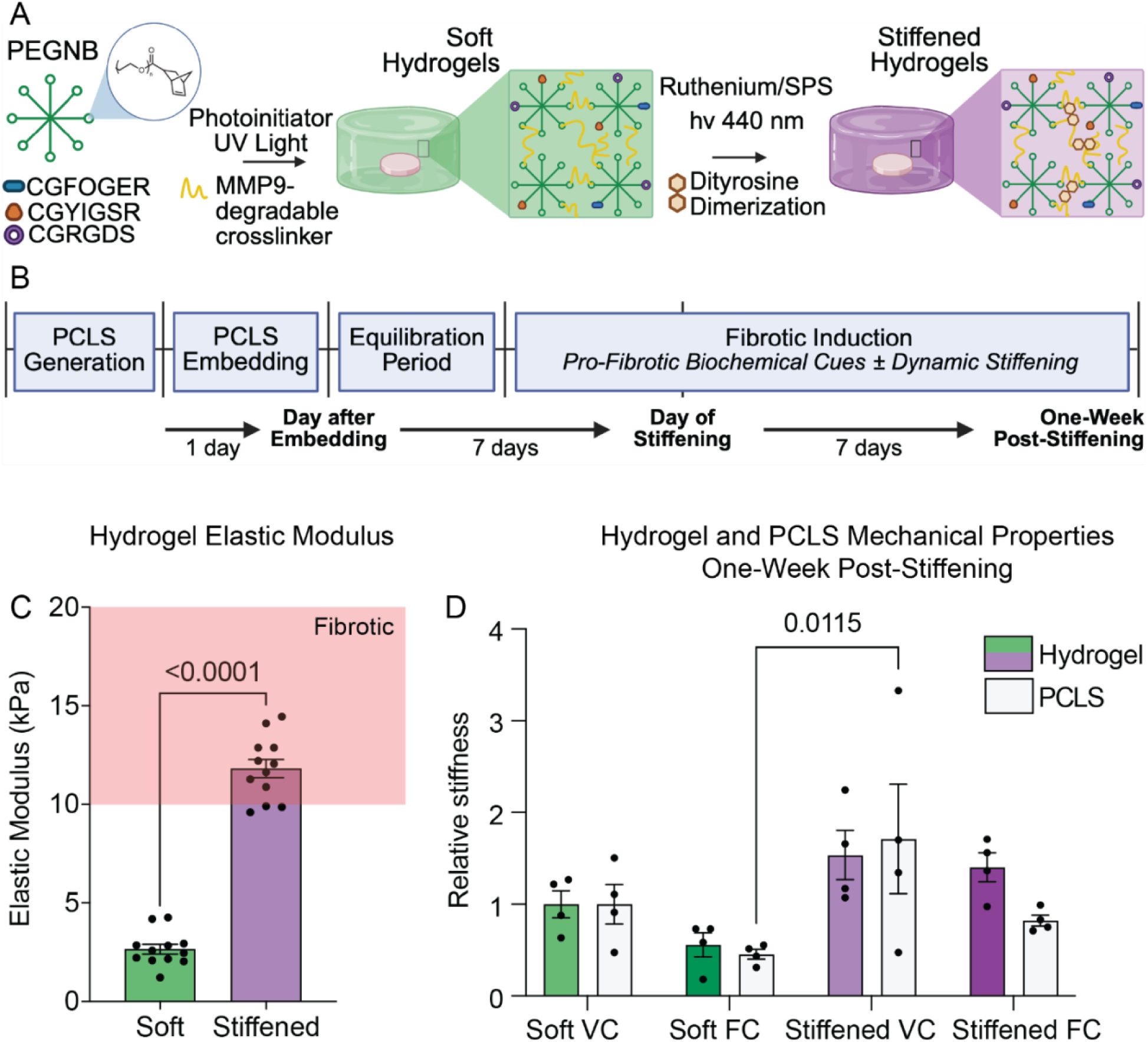
An extended culture hydrogel-embedded PCLS model recapitulates features of early pulmonary fibrosis. (A) PCLS were embedded in a soft PEGNB hydrogel formulation containing a MMP9-degradable crosslinker sequence, CGFOGER peptide, CGYIGSR peptide, and CGRGDS peptide one-day after slicing. Samples intended to be stiffened were incubated with ruthenium and SPS for 6 h one week after embedding and then underwent 5 min of visible blue light (hν 440nm) exposure that dynamically stiffened the formulation because of additional dityrosine dimerization. (B) Schematic representation of the timeline used for cell culture experiments that highlights outputs, dynamic stiffening, and pro-fibrotic biochemical cue exposures. (C) Parallel plate rheology with acellular PEGNB hydrogels showed the average elastic modulus (E) of soft hydrogels (E = 2.65 ± 0.25 kPa) fell within the range of healthy lung tissue (1-5 kPa), while the average elastic modulus of stiffened hydrogels (E = 11.81 ± 0.46 kPa) increased into the range of fibrotic tissue (>10 kPa) (pink shaded zone). Columns represent mean ± SEM, n = 12. Symbols represent technical replicates. Statistical significance was determined by an unpaired t-test. (D) The elastic modulus of hydrogel and tissue regions within hydrogel-embedded PCLS samples was measured by atomic force microscopy. Relative stiffness values are presented using a baseline correction against the hydrogel and tissue soft VC averages. Columns represent mean ± SEM, n = 4. Symbols represent the average baseline corrected relative elastic modulus from each biological replicate. Statistical significance was assessed using two-way ANOVA with Tukey’s multiple comparisons test.

Additionally, since two batches of PEGNB were used in these studies and their functionalization varied slightly, a small range of PEGNB concentrations (3.75 wt% and 4 wt%) were tested to ensure the elastic modulus value targets matched. There were no statistical differences in the elastic modulus values between the 3.75 wt% and 4 wt% PEGNB batches of hydrogels based on rheological measurements (**Supplemental Figure 2**). This same principal was then applied to PCLS-embedded constructs, since the tissue itself also contained tyrosine residues via the native extracellular proteins.

Standard protocols generated mouse PCLS. Briefly, mouse lungs were filled with 1.5 wt% agarose and sliced on a vibratome. The 500-μm tissue slices were then cut into 4-mm samples, incubated overnight to help facilitate agarose removal, and embedded within the engineered soft PEGNB hydrogels. Custom silicone molds with 8-mm diameter wells produced hydrogel-embedded PCLS as previously described (26, 27). The thiol-to-norbornene ratio of 0.7 was selected so free reactive groups would remain after the bottom layer of hydrogel was polymerized. Free norbornenes allowed for the top layer of hydrogel mixture to crosslink with the biomaterial underneath and produce a solid hydrogel structure encapsulating the PCLS tissue.

Within fibrotic lung tissue, increases in inflammatory mediators and stiffness are observed as the disease progresses (29, 47, 59, 60). To recapitulate these features of fibrosis progression *ex vivo*, hydrogel-embedded PCLS from control (non-diseased) mouse lungs were exposed to pro-fibrotic biochemical cues and/or dynamic microenvironmental stiffening. A fibrosis cocktail (5 ng/ml TGF-β, 10 ng/ml PDGF-AB, and 5 μM LPA) was supplemented into the hydrogel-embedded PCLS medium every 4 days, to expose a portion of the samples tested to pro-fibrotic biochemical cues. Likewise, a subset of constructs were dynamically stiffened to achieve an elastic modulus representative of fibrotic lung tissue (>10 kPa) (28, 30). Ruthenium and SPS photoinitiators were swollen into the constructs for 6 h and exposed to visible blue light (440 nm) on day 8. Initially, we expected that the cells present within the PCLS constructs would cause enough remodeling through enzymatic degradation or new ECM deposition that the elastic modulus of the embedding hydrogel would change over time. Therefore, to test this hypothesis, supernatant was collected from cell-laden hydrogel-embedded PCLS samples and added to soft acellular hydrogels every 48 h. It was expected that this supernatant would contain matrix-remodeling enzymes secreted by hydrogel-embedded PCLS and that adding it to acellular hydrogels would enable analyses of how these secreted factors influenced the biomaterial stiffness. Although not significant, a slight decrease in the elastic modulus of acellular hydrogels was measured after 15 days when dosed with the soft FC, stiffened VC, and stiffened FC hydrogel-embedded PCLS supernatants (**Supplemental Figure 4**). These findings motivated more extensive mechanical characterization of embedding hydrogels and PCLS tissue over time using AFM.

AFM was performed on the constructs to map spatial variations in stiffness across both the embedding hydrogel and PCLS tissues, each of which can be dynamically stiffened through dityrosine dimerization (14) (**Figure 1D)**. By day 15, stiffened hydrogels and tissues exhibited greater relative stiffness than the soft counterparts. Overall, stiffening and/or biochemical cue exposure produced wide variability in stiffness within each condition, which reflects the heterogenous nature of tissue remodeling in response to fibrotic stimuli and recapitulates heterogeneity observed in vivo (**Figure 1D).** These findings together suggested that the cells within hydrogel-embedded PCLS were actively remodeling the microenvironment in response to pro-fibrotic biochemical cues and matrix stiffening.

### Hydrogel-embedded PCLS maintained long-term cellular viability

Cellular viability was assessed in hydrogel-embedded PCLS by live/dead staining on days 1, 8, and 15. Across all conditions, cell viability remained high (>80%) throughout the culture period. Representative fluorescent images illustrating total cells identified by nuclei (blue) and dead cells (green) within soft and dynamically stiffened constructs at one-week post-stiffening (day 15) under the FC conditions are shown in **Figure 2A**.

**Figure 2.**
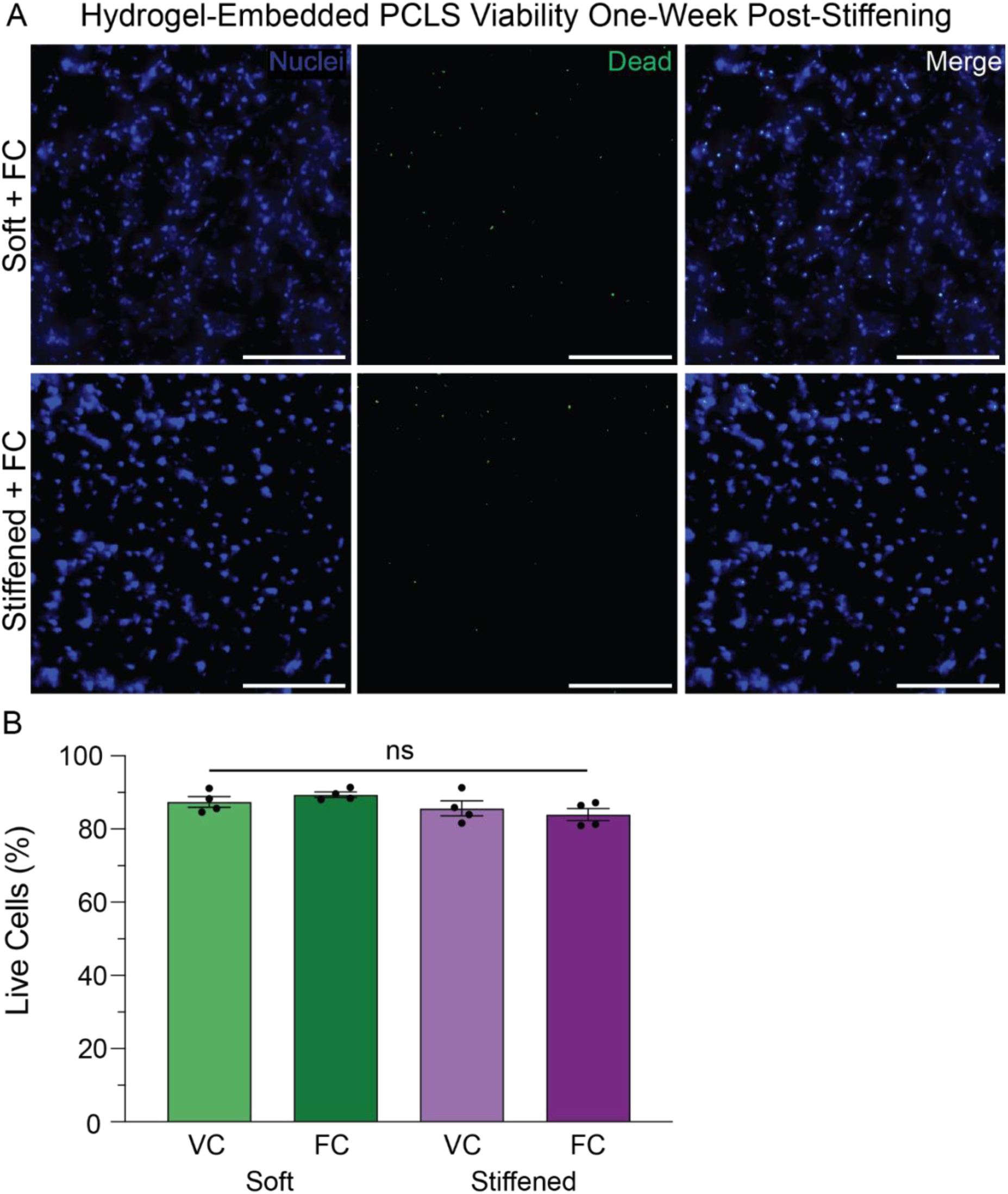
Dynamic stiffening and exposure to pro-fibrotic biochemical cues did not induce overt toxicity or significantly alter cellular viability in hydrogel-embedded PCLS. (A) Hydrogel-embedded PCLS were stained for 1 h using a commercial viability kit which showed total cell nuclei (blue) and dead cell (green) staining. Representative maximum-intensity z-stack projections are shown of fibrosis cocktail (FC)-exposed soft and dynamically stiffened hydrogel-embedded PCLS constructs one week after stiffening (day 15). Scale bar, 100 μm. (B) Viability was quantified by comparing the number of dead cells to the total number of cell nuclei. Results showed that the average PCLS viability remained above 80% across all conditions at one-week post-stiffening (day 15). No statistically significant differences were detected between VC- and FC-treated samples within either the soft or stiffened hydrogel groups. Statistical significance was assessed using two-way ANOVA with Tukey’s multiple comparisons test. Bars represent mean ± SEM; n = 4. Symbols indicate individual experimental replicates (average of 3-6 individual samples from separate embedding rounds).

One day later (on day 1) embedding, the average percentage of live cells within hydrogel-embedded PCLS was 89% (**Supplemental Figure 3A**). On the day of stiffening (day 8), average viability was 90% in the soft VC condition, 93% in the soft FC condition, and 88% in both the stiffened conditions (**Supplemental Figure 3B**). One week post-stiffening (day 15), average viability remained high with 87% live cells in the soft VC condition, 89% in the soft FC condition, 86% in the stiffened VC condition, and 84% in the stiffened FC condition (**Figure 2B**). Although a modest reduction in viability was observed in stiffened constructs relative to soft hydrogels, no statistically significant differences were detected among conditions at any time point.

### Pro-fibrotic biochemical cue exposure and dynamic stiffening recapitulated fibrotic matrix deposition and gene expression patterns in hydrogel-embedded PCLS

To assess how hydrogel-embedded PCLS responded to pro-fibrotic stimulation, fluorescent imaging visualized newly deposited (nascent) ECM proteins within the constructs. Representative images from all four experimental conditions on day 15 are shown in **Figure 3A**, with nascent proteins (green), cellular cytoplasm (red), and cell nuclei (blue) visualized by fluorescence. Quantification of these images revealed that the stiffened and FC-exposed condition produced the highest level of nascent protein deposition (**Figure 3B**). Significant increases were observed when comparing VC soft samples at the day of stiffening (day 8) to VC soft samples one week later (day 15) (p = 0.0153). A similar pattern was detected in VC stiffened samples (p < 0.001) and in FC stiffened samples (p = 0.0092). Within the FC-treated group, stiffened constructs demonstrated greater nascent ECM deposition than soft constructs at day 15 (p = 0.0392). Finally, comparison of the least fibrotic condition (soft VC, day 8) to the most fibrotic condition (stiffened FC, day 15) demonstrated a highly significant increase in nascent ECM deposition (p < 0.0001). This comparison establishes the dynamic range of the model, demonstrating that increased matrix stiffness, fibrotic biochemical exposure, and extended culture promote increased nascent ECM deposition consistent with fibrotic progression.

**Figure 3.**
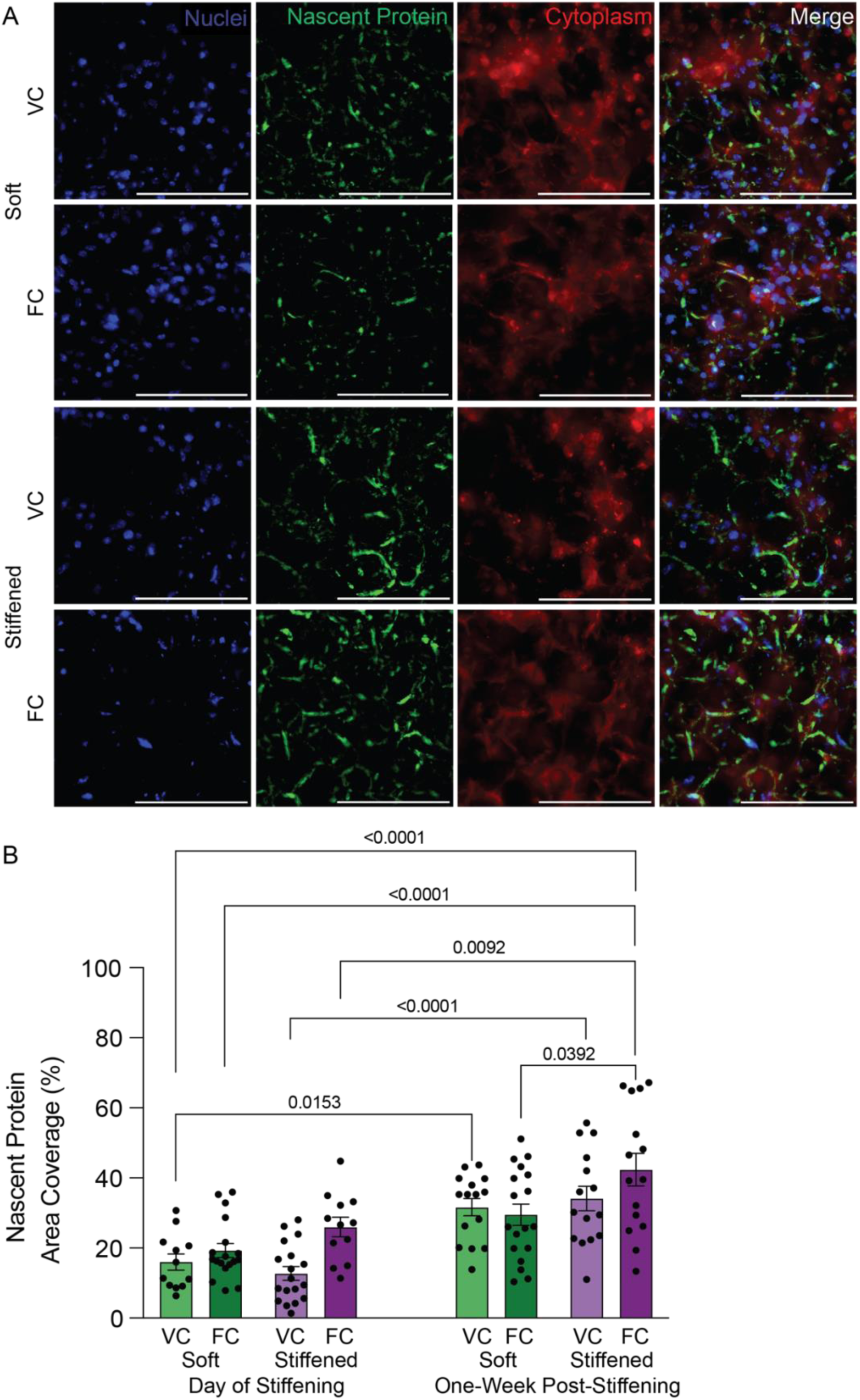
Stiffening and pro-fibrotic biochemical cue exposure increased nascent protein deposition in hydrogel-embedded PCLS. (A) Hydrogel-embedded PCLS samples were fluorescently stained for cell nuclei (blue), nascent ECM proteins (green), and cellular cytoplasm (red) on day 15. Images shown are representative maximum-intensity z-stack projections from each condition. Scale bar, 100 μm. (B) Nascent ECM protein was quantified as the percentage of fluorescent intensity area coverage present within the FITC channel from the maximum-intensity z-stack projection images. Statistical significance was assessed using a two-way ANOVA test with Tukey’s test for multiple comparisons. Bars represent mean ± SEM; n = 12-18. Symbols denote technical replicates.

To evaluate epithelial responses to combined pro-fibrotic biochemical stimulation and dynamic stiffening, mRNA was isolated from hydrogel-embedded PCLS across multiple experimental conditions and time points. RT-qPCR was performed to assess gene expression associated with ATII identity (**Figure 4A**), ATII-to-ATI transitional states (**Figure 4B**), and mature ATI epithelium (**Figure 4C**) (61–63). Based on previous studies, increased microenvironmental stiffness and FC exposure were expected to suppress ATII markers while promoting ATII-to-ATI transitional and ATI gene expression (16–18, 20, 25). Comparisons between day 8 (day of stiffening) and day 15 (one-week post-stiffening) revealed several notable changes. *Sftpc* expression increased in FC stiffened samples (p = 0.0125) (**Figure 4A**). *Krt8* expression was upregulated in both VC (p = 0.0248) and FC (p = 0.0420) stiffened samples (**Figure 4B**). Similarly, *Hopx* expression increased in both VC (p = 0.0013) and FC (p = 0.0003) stiffened groups (**Figure 4C**). By day 15, additional increases were detected when comparing VC soft to VC stiffened samples (p = 0.0065), FC soft to FC stiffened samples (p = 0.0159), and VC soft to FC stiffened samples (p = 0.0026) (**Figure 4C**). In contrast, only one significant decrease was observed: Ager expression declined from day 8 to day 15 in the FC soft group (p = 0.0463) (**Figure 4C**).

**Figure 4.**
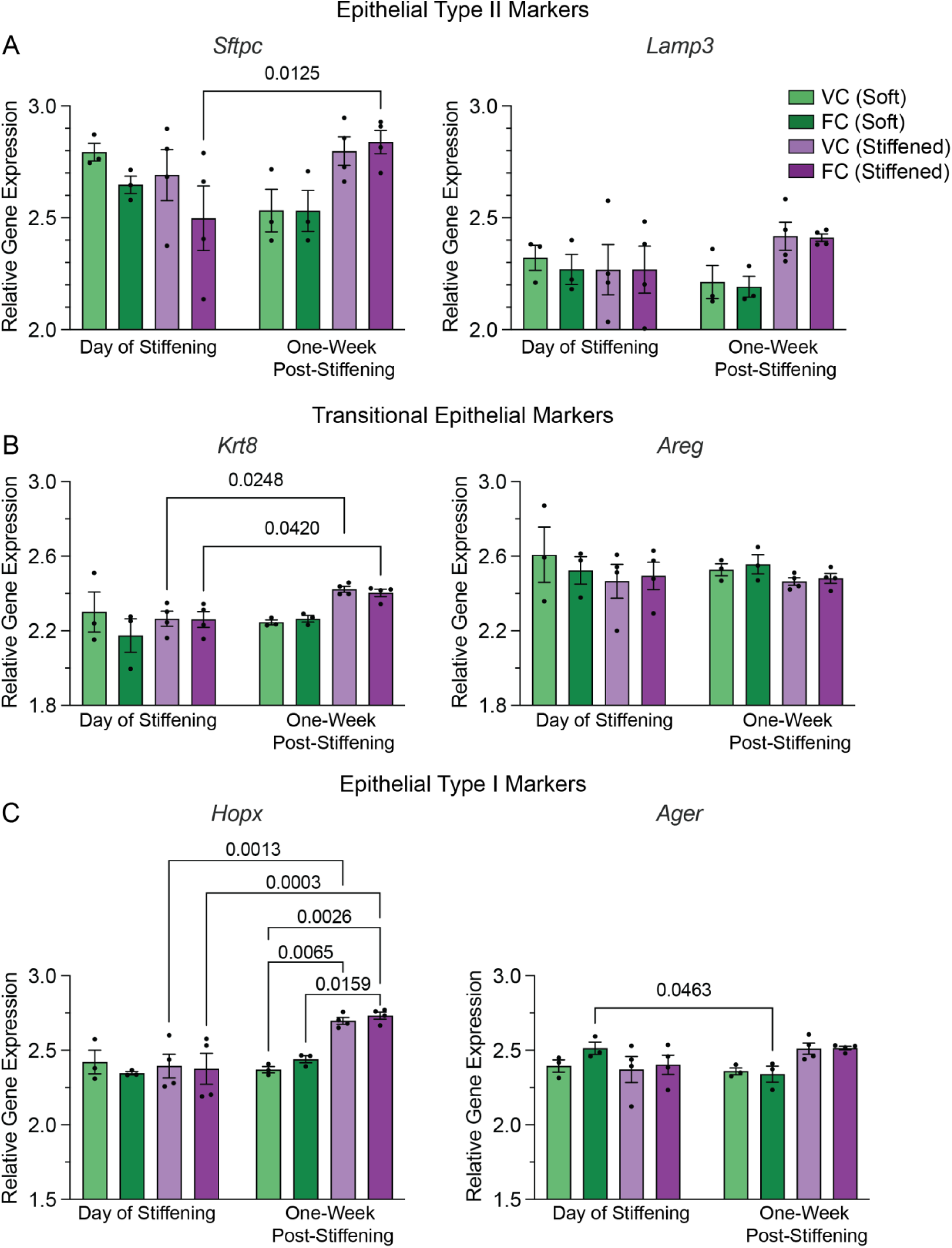
Stiffening and exposure to pro-fibrotic biochemical cues altered epithelial cell gene expression patterns in hydrogel-embedded PCLS. (A) RNA was isolated from PCLS samples at two timepoints, on the day of stiffening and at one-week post-stiffening. To assess ATII gene expression, *Sftpc* and *Lamp3* were chosen for RT-qPCR analysis. (B) To assess transitional epithelial cell gene expression, *Krt8* and *Areg* were chosen for RT-qPCR analysis. (C) To assess ATI gene expression, *Hopx* and *Ager* were chosen for RT-qPCR analysis. Statistical significance for all analyses was determined using two-way ANOVA with Tukey’s multiple comparisons test. Bars represent mean ± SEM; n = 3-4. Symbols denote experimental replicates (pooled PCLS from each separate embedding round)

Overall, while a broad downregulation of epithelial gene expression was anticipated, the observed increases, particularly among transitional and ATI-associated markers, suggest that hydrogel-embedded PCLS may preserve epithelial activation and repair responses observed *in vivo* under *ex vivo* fibrotic stimulation (26, 64–66).

In contrast to the variable epithelial signatures, fibroblast activation markers, including *Col1a1*, *Cthrc1*, *Ltbp2*, and *Fn1*, displayed more consistent expression patterns across experimental conditions, except for *Ctgf*, which diverged from these trends (**Figure 5A**). As anticipated, increased microenvironmental stiffness and FC exposure generally promoted fibroblast activation (29, 30, 67, 68). *Ltbp2* expression increased significantly between VC soft and VC stiffened samples (p = 0.0032), FC soft and FC stiffened samples (p = 0.0035), and VC soft and FC stiffened samples at day 8 (p = 0.0014) (**Figure 5A**). Additional increases were found when comparing day 8 to day 15 within both VC stiffened (p = 0.0202) and FC stiffened (p = 0.0057) groups, as well as between VC soft day 8 and VC stiffened day 15 (p < 0.0001) and FC soft day 8 and FC stiffened day 15 (p = 0.0002) (**Figure 5A**). In contrast, *Ctgf* expression decreased significantly in FC stiffened constructs from day 8 to day 15 (p = 0.0227).

**Figure 5.**
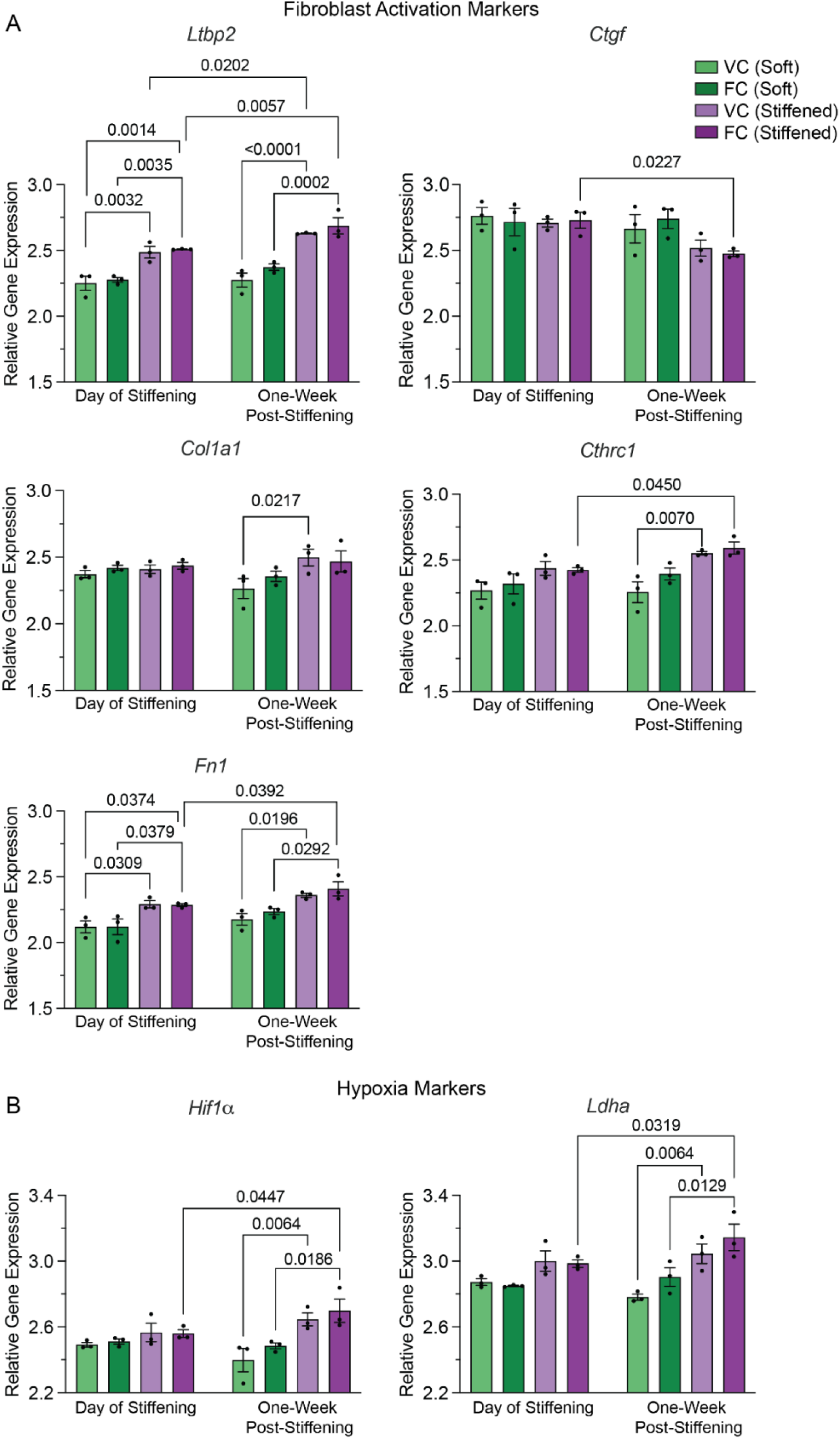
Fibroblast activation and hypoxia-associated gene expression were modulated by microenvironmental stiffness and pro-fibrotic biochemical cues. (A) RNA was isolated from PCLS samples at two timepoints, on the day of stiffening and at one-week post-stiffening. To assess fibroblast activation gene expression at these timepoints, *Col1a1*, *Cthrc1*, *Ltbp2*, *Fn1*, and *Ctgf* were chosen for RT-qPCR analysis. (B) To assess hypoxia gene expression within the PCLS samples at these timepoints, *Hif1α* and *Ldha* were chosen for RT-qPCR analysis. Statistical significance for all analyses was determined using two-way ANOVA with Tukey’s multiple comparisons test. Bars represent mean ± SEM; n = 3. Symbols denote experimental replicates (pooled PCLS from each separate embedding round).

For *Col1a1*, a significant increase was observed between VC soft and VC stiffened samples on day 15 (one-week post-stiffening) (p = 0.0217) (**Figure 5A**). *Cthrc1* expression showed a similar pattern (p = 0.007), in addition to an increase between FC stiffened samples at day 8 (day of stiffening) and day 15 (p = 0.045) (**Figure 5A**). *Fn1* expression also increased across multiple comparisons, including VC soft vs. VC stiffened (day 8; p = 0.0309), FC soft vs. FC stiffened (day 8; p = 0.0379), VC soft vs. FC stiffened (day 8; p = 0.0374), FC stiffened day 8 vs. day 15 (p = 0.0392), VC soft vs. VC stiffened on day 15 (p = 0.0196), and FC soft vs. FC stiffened on day 15 (p = 0.0292) (**Figure 5A**).

As hypoxia is a known contributor to the fibrotic progression (69–72), both *Hif1α* and *Ldha* were also evaluated. Both genes exhibited notable increases in FC stiffened samples between day 8 and day 15 (*Hif1α*: p = 0.0447; *Ldha*: p = 0.0319). Significant increases were additionally observed between VC soft and VC stiffened samples on day 15 (*Hif1α*: p = 0.0064; *Ldha*: p = 0.0064) and between FC soft and FC stiffened samples on day 15 (*Hif1α*: p = 0.0186; *Ldha*: p = 0.0129) (**Figure 5B**). Across fibroblast activation and hypoxia-associated genes, FC stiffened constructs at one week post-stiffening consistently exhibited the highest relative expression.

Collectively, these data demonstrate that both pro-fibrotic biochemical cues and increased stiffness strongly influence fibroblast and hypoxia-associated gene expression within hydrogel-embedded PCLS, complementing the epithelial responses described above. The emergence of clearer gene expression patterns at later time points suggests that extended culture may enable researchers to further elucidate mechanisms underlying fibrotic activation and progression.

### Hydrogel-embedded PCLS responded to anti-fibrotic drug treatment

To further assess the responsiveness of the hydrogel-embedded PCLS model to potential anti-fibrotic therapy, we selected Nintedanib as the first drug candidate. Since fibrotic induction was observed by day 15 in the earlier experiments, Nintedanib was administered from one week post-stiffening (day 15) to two weeks post-stiffening (day 22) to evaluate whether the drug could mitigate the ongoing fibrotic response and slow progression (**Figure 6**). Stiffened hydrogel-embedded PCLS received either FC exposure and Nintedanib treatment together, or FC exposure alone as a control. As shown in **Supplemental Figure 5**, cell viability remained high throughout the extended culture period and Nintedanib exposure, consistent with prior live/dead assessments. On day 22, viability averaged 87% in stiffened FC samples and 88% within the stiffened FC samples treated with Nintedanib (**Supplemental Figure 5B**). Nintedanib treatment resulted in modest, non-significant downregulation of all epithelial markers (**Figure 6A**). Similarly, all fibroblast activation markers displayed slight decreases following Nintedanib treatment, except for *Ltbp2* (**Figure 6B**). Despite the range of responses, slices pooled from the same experimental replicate tended to follow the same patterns, as seen visually by using different shapes for the individual dots (**Figure 6**).

**Figure 6.**
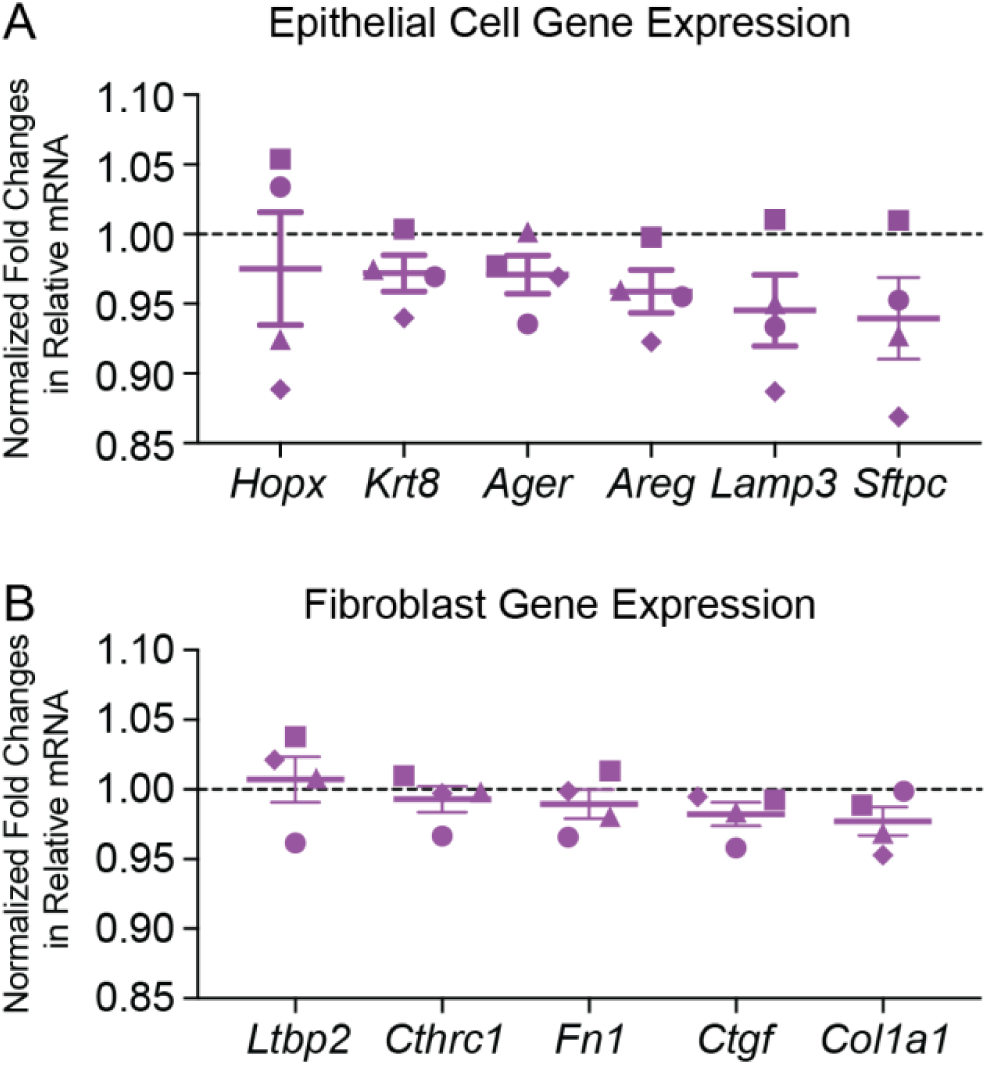
Embedded-PCLS lung models showed modest responsiveness to anti-fibrotic treatment after 3 weeks in culture. Nintedanib was administered to stiffened hydrogel-embedded PCLS from days 16 to day 22 (one to two weeks post-stiffening) to assess its ability to mitigate ongoing fibrotic progression. (A) Relative gene expression of epithelial cell markers in stiffened hydrogels following Nintedanib treatment on day 22, normalized to the FC-only controls. (B) Relative gene expression of fibroblast activation markers in stiffened hydrogels following Nintedanib treatment on day 22 (two-weeks post-stiffening), normalized to the FC-only controls. All statistical analyses assessed significance by unpaired t-tests with Welch comparison. Bars represent mean ± SEM, n=4. Symbols indicate experimental replicates (pooled hydrogel-embedded PCLS samples from each embedding round).

## Discussion

This work integrates several established approaches to create an extended-culture PCLS model of pulmonary fibrosis using engineered PEGNB hydrogels with tunable mechanical properties. Healthy (non-diseased) mouse PCLS were embedded within hydrogels and exposed to pro-fibrotic biochemical cues and/or dynamic stiffening to recapitulate key aspects of fibrosis progression under controlled conditions. This platform supported multi-week culture while maintaining high cellular viability, consistent with previous reports demonstrating that hydrogel embedding preserves cell viability and function (26, 27). Live/dead analyses showed over 80% viability across three weeks of culture in all tested conditions. To emulate repetitive injury models used *in vivo*, including repetitive bleomycin injury (48, 73), a FC was administered every four days. A subset of hydrogel-embedded PCLS also underwent dynamic stiffening to model the progressive mechanical changes associated with fibrotic lung disease.

Lung fibrosis is characterized by alterations in both ECM composition and matrix stiffness (29, 40, 47, 58, 60), along with epithelial cell dysfunction, fibroblast activation, and the accumulation of excess ECM (60, 74). Repeated epithelial injury promotes fibroblast activation through secretion of mediators, like TGF-β (75–78), which increases oxygen consumption and shifts fibroblast metabolism toward aerobic glycolysis (79–82). Consistent with these established pathways, hypoxia-associated gene expression was highest in the FC-treated, stiffened condition at one week post-stiffening. Unlike several previous PCLS studies reporting rapid epithelial loss and significant cell death (16, 18, 83–85), epithelial cells in hydrogel-embedded PCLS remained present across all conditions over multiple weeks. This sustained epithelial presence enables temporal analysis of epithelial behavior under physiologically relevant stimulation by biochemical and mechanical cues. The observed trends suggest ongoing epithelial renewal and differentiation and indicate that the cells tolerated repeated FC exposure more effectively than reported in some earlier studies (16, 17, 20). Future work incorporating cell-type-specific staining and/or lineage tracing (86–90) would clarify subpopulation dynamics in greater detail for mouse or human hydrogel-embedded PCLS.

Given the central role of cell-matrix interactions in fibrosis, it remains a subject of investigation whether biochemical cues or mechanical properties are the more potent drivers of disease progression (28–30, 60, 74, 91, 92). RT-qPCR analysis in this model indicated that the matrix had a dominant influence on fibrotic gene expression (28, 29, 47), although biochemical cues and extended time in culture also produced significant changes. Several significant differences emerged when comparing soft versus stiffened hydrogel-embedded PCLS and early versus later timepoints. Markers affected by these variables included *Sftpc*, *Krt8*, *Hopx*, *Ager*, *Cthrc1*, *Ctgf*, *Ltbp2*, *Fn1*, *Hif1α*, and *Ldha*. When experimental conditions were arranged from lowest-expected (soft, VC-exposed) to highest-expected fibrotic (stiffened, FC-exposed) severity, relative gene expression followed the anticipated pattern most clearly at day 15, indicating that the combination of increased stiffness and biochemical stimulation produced the most robust fibrotic activation after two weeks of induction.

Therapeutic responsiveness was evaluated using the FDA-approved anti-fibrotic drug Nintedanib at a commonly used 10 μM concentration (16, 17). Although Nintedanib has been shown to reduce epithelial cell injury (20) and inhibit fibroblast activation (3, 17, 93), only modest reductions in epithelial and fibroblast activation markers were observed in this study. Since treatment was applied exclusively to stiffened constructs (conditions already exhibiting peak activation after two weeks of induction), the dynamic range for therapeutic reversal may have been limited. Alternative treatment windows, earlier intervention, or longer time periods may produce stronger responses in future studies. Evaluation of additional anti-fibrotics, including the newly FDA-approved anti-fibrotic drug Nerandomilast (94–96), would further establish the suitability of hydrogel-embedded PCLS models as a new approach methodology for drug testing.

Hydrogel-embedded PCLS models may decrease the gap between benchtop research and clinical translation, as well as advance anti-fibrotic drug screening in the future. However, several system-specific considerations should be acknowledged. Mouse PCLS were used due to limited access to human tissue. Repeating this work with human PCLS is essential, particularly because fibrosis has been shown to resolve spontaneously in young mice that were challenged with one bleomycin instillation, but not in humans (48, 97–99). The hydrogel was designed to approximate healthy lung stiffness at baseline and then stiffen to match fibrotic lung tissue stiffness after 8 days in culture. This was observed in the stiffened VC hydrogel and tissue constructs measured by AFM, which remained stiffer than the soft samples at day 15. The inclusion of the CGYIGSR peptide and MMP9-degradable crosslinker enabled secondary stiffening via tyrosine-mediated photochemical reactions, which was verified in acellular hydrogels. Additionally, although fibrotic activation was induced over the first two weeks in these initial studies and anti-fibrotic treatment was assessed in week three, prior work indicates that hydrogel-embedded PCLS could have likely been maintained for substantially longer (up to six weeks) (26, 27, 29).

Preliminary enzymatic degradation studies were performed by culturing acellular hydrogels in the supernatant collected from the different cell-laden hydrogel-embedded PCLS constructs. Although modest and non-significant, subtle decreases in the elastic modulus of these acellular hydrogels was observed when cultured using the soft FC, stiffened VC, and stiffened FC supernatants. Additionally, nascent ECM deposition was greatest in stiffened, FC-exposed samples. AFM results further demonstrated that within soft hydrogels, FC exposure resulted in decreased relative stiffness by day 15. Interestingly, even under stiffened conditions, FC-exposed samples also exhibited lower relative stiffness compared to stiffened VC samples. Together, these findings suggest that FC exposure promotes a more pronounced remodeling response, potentially characterized by increased matrix turnover and structural destabilization, which aligns with previous findings (100–102). Transient mechanisms such as elevated MMP activity, basement membrane disruption, or loss of alveolar structural integrity could contribute to reduced stiffness measurements prior to substantial collagen accumulation and crosslinking (103, 104). Such biphasic mechanical behavior – initial softening followed by progressive stiffening – has been described in injury-driven fibrotic models and may indicate that this system captures early, degradation-dominant remodeling processes (101, 102). The use of genetically modified reporter mice would facilitate identification of the cellular sources of the newly synthesized ECM proteins that were observed in these studies (54).

Additional adjustment of the hydrogel formulation may improve mechanotransductive specificity. Removing mimetic peptides or replacing the linear CGFOGER, which has mixed reported efficacy for supporting integrin binding (105, 106), with a helical analog may reduce confounding variables. Testing hydrogels without the MMP9-degradable crosslinker, so that no tyrosine residues are present within the embedding hydrogel formulation itself, could facilitate evaluation of which tyrosines are actively participating in the stiffening mechanism. Further characterization of secreted proteases including MMPs, could also clarify the extent of matrix remodeling over time. Another limitation of this work is the use of collagen I (*Col1a1*) gene expression as a primary marker of fibroblast activation (68). Although *Col1a1* is widely used to indicate matrix-producing fibroblasts, it is not cell-type specific and can also be upregulated in epithelial cells undergoing epithelial-mesenchymal transition (EMT). In this study, *Col1a1* is interpreted as a fibroblast activation marker due to its established relevance in lung fibrosis; however, this approach does not distinguish between mesenchymal and epithelial sources of expression. Notably, single-cell transcriptomic studies (68, 87, 107) demonstrate that *Col1a1* expression is predominantly enriched in fibroblast and myofibroblast populations, with epithelial expression typically limited to transitional states and lower in magnitude. Nevertheless, the contribution from EMT-associated epithelial cells should be further explored and decoupled in the future.

This biomaterial-based PCLS platform is well positioned for integration into emerging analytical and engineering approaches. Combining hydrogel-embedded PCLS with transcriptomic, proteomic, and high-resolution imaging strategies deepen mechanistic insight into ECM-driven regulation of cell behavior, as demonstrated in recent studies applying sequencing approaches to engineered PCLS systems (23). Continued innovations in biomaterial design and microfluidic technologies are expected to expand the utility of PCLS for modeling complex lung diseases and evaluating personalized therapeutic responses.

In conclusion, *ex vivo* PCLS models of pulmonary fibrosis have traditionally relied on static, non-embedded, short-term culture systems. The approach presented here demonstrates the feasibility of embedding PCLS in engineered hydrogels capable of delivering dynamic mechanical cues and supporting multi-week culture. Across viability, gene expression, and nascent ECM deposition analyses, the findings indicate that this system recapitulates key features of early fibrotic activation and offers an improved platform for studying long-term fibrosis progression and evaluating anti-fibrotic therapeutics.

## Data Availability Statement

The data that support the findings of this study are openly available in Mendeley Data at doi:10.17632/szrfjhhj3n.2.

## Acknowledgements

Portions of Figure 1 (Magin, C. (2025), https://BioRender.com/734k3a0) were created using BioRender. The authors sincerely thank Dr. Andrei Krivoi (University of Colorado Anschutz) for his time and assistance in teaching the members of the Magin laboratory a variety of precision-cut lung slicing techniques. Additionally, we would like to thank Anton D. Kary (University of Colorado Denver | Anschutz) for contributing text to an early draft of the introduction. We also are grateful to Dr. David Schwartz (University of Colorado Anschutz) and the Division of Pulmonary, Allergy, and Critical Care Medicine for access to shared equipment, specifically the vibratome that produced PCLS for a number of these experiments. We would also like to acknowledge Hans J. Kaper (University of Groningen) and Prashant K. Sharma (University of Groningen) for teaching the technique of low load compression testing, Rocío Fuentes Mateos (University of Groningen) and Reinoud Gosens (University of Groningen) for their assistance in procuring mouse lung tissue, and Manon E. Woest (University of Groningen) for her technical assistance in initial PCLS generation. Even though this work was not included in the final manuscript, we deeply appreciate the time and effort they devoted to these preliminary studies.

This work was supported by funding from the National Heart, Lung, and Blood Institute of the National Institutes of Health (NIH) under awards R01 HL153096 (CMM, RB, AET, HN) and T32 HL072738 (AET), the Gates Summer Internship Program (TY), the NIH/NCATS Colorado CTSA Grant Number T32 TR004367 and UM1 TR004399 (HN), and through the European Respiratory Society Short Term Research Fellowship Program under STRTF202410-01192 (AET).

