## Supplemental Material for "Hydrogel-Embedded Precision-Cut Lung Slices Recapitulate Fibrotic Gene Expression and Enable Therapeutic Response Evaluation"

### Supplemental Methods

*Supernatant Studies*

Soft acellular hydrogels were fabricated as described in the main text, equilibrated in PCLS medium, and maintained at 37°C and 5% $CO_2$. Every 48 hours, the medium was replaced with either fresh PCLS medium (control) or conditioned medium collected from hydrogel-embedded PCLS from the following conditions: soft VC, soft FC, stiffened VC, or stiffened FC samples. For conditioned groups, 300 μL of supernatant from a specific hydrogel-embedded PCLS well was transferred to the matched acellular hydrogels at each media change, ensuring that conditioned medium from the same source well was consistently applied across timepoints. Then, the acellular hydrogels were measured using parallel plate rheology in accordance with the methods found within the manuscript.

### Supplemental Results

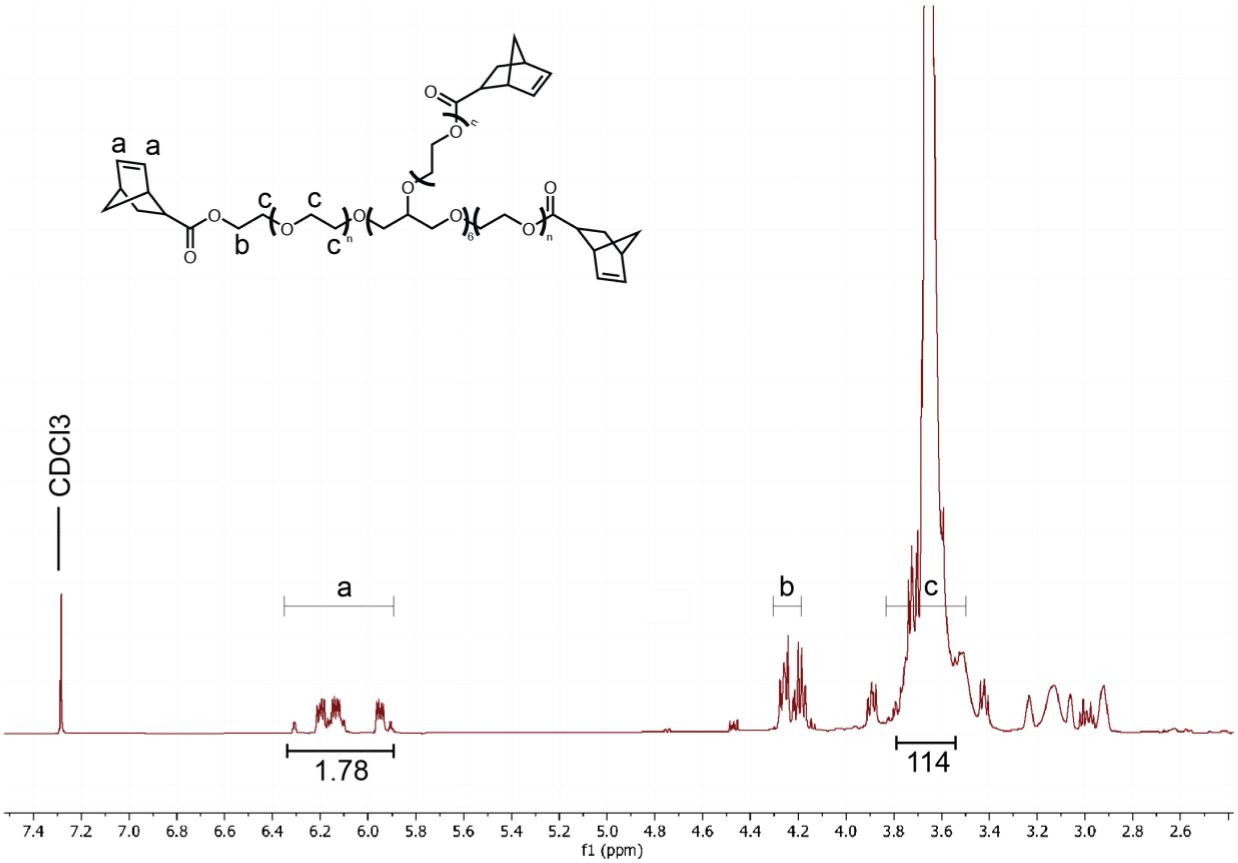


**Supplemental Figure 1.** $^1$H NMR results of 10 kg/mol PEGNB (Bruker DPX-400 FT NMR spectrometer). δH (ppm) (300 MHz, $CDCl_3$): 3.71 (s, 114H, PEG $CH_2$-$CH_2$), 4.1-4.2 (m, 2H, -$CH_2$-O), 5.9-6.2 (m, 2H, –CH=CH-). NMR shows an 89% quantitative norbornene functionalization based on a comparison of the alkene protons from norbornene to the theoretically expected number of alkene protons, using the ethylene glycol protons as a reference.

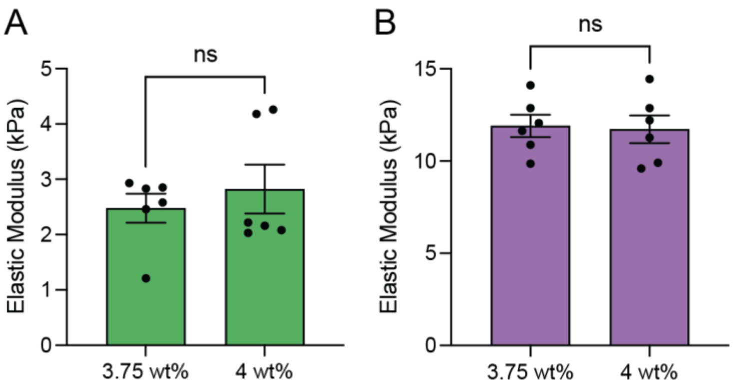


**Supplemental Figure 2.** Additional mechanical characterizations of samples. (A) Using rheological characterization, there were no statistical differences in the measured elastic modulus within soft hydrogels formed using either a 3.75 wt% or 4 wt% PEGNB formulation. (B) There were no statistical differences in the elastic modulus measurements within the stiffened hydrogels that were formed using either a 3.75 wt% or 4 wt% initially soft formulation based on rheological characterization. Columns represent mean ± SEM, n=6 (ns = no significance). Symbols represent technical replicates. All statistical analyses determined significance by an unpaired t-test with Welch comparison.

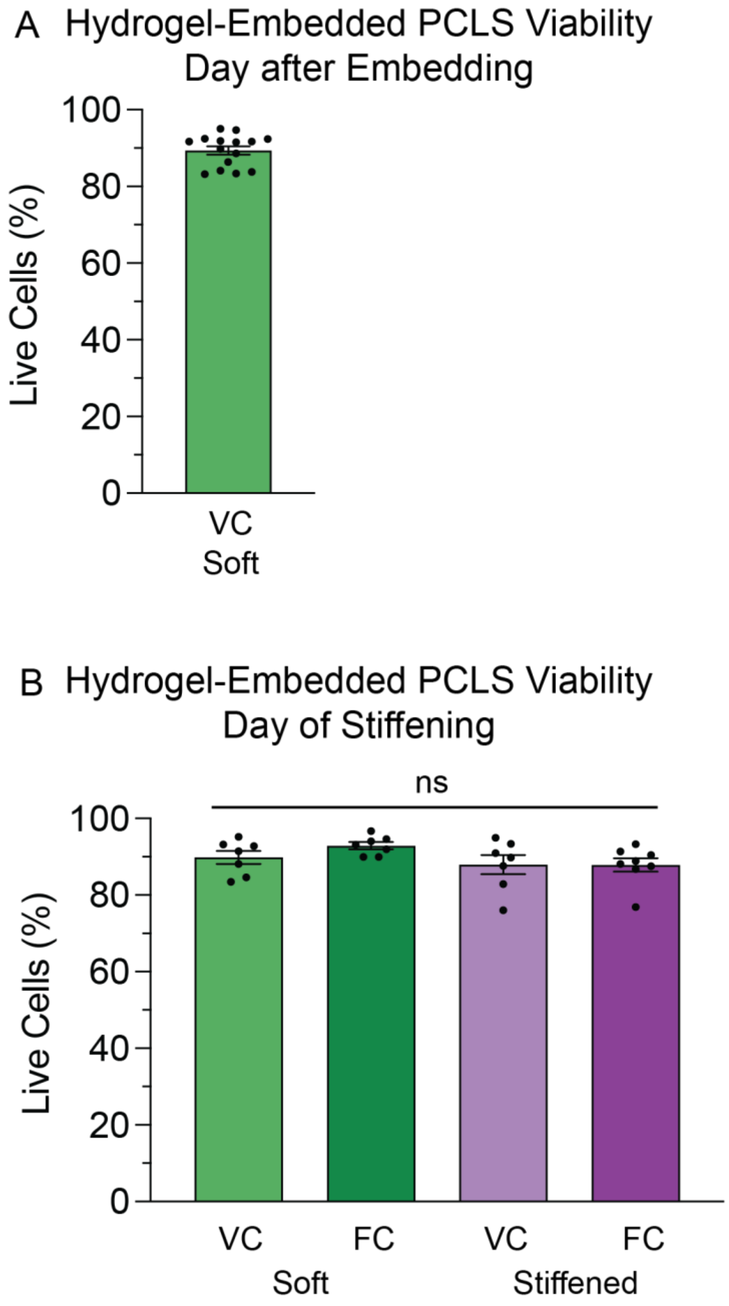


**Supplemental Figure 3.** Quantification of live/dead images support that PCLS embedding and dynamic stiffening did not negatively impact viability. A) Approximately 90% of cells remained viable one day post PCLS embedding. B) After one week in culture, PCLS viability remained similar to the initial cell viability. There were no statistical differences across sample types on the day of stiffening. Columns represent mean ± SEM, n=7-8. Symbols represent technical replicates, and statistical significance was determined by an ordinary one-way ANOVA test with Tukey’s multiple comparisons test, ns = not significant.

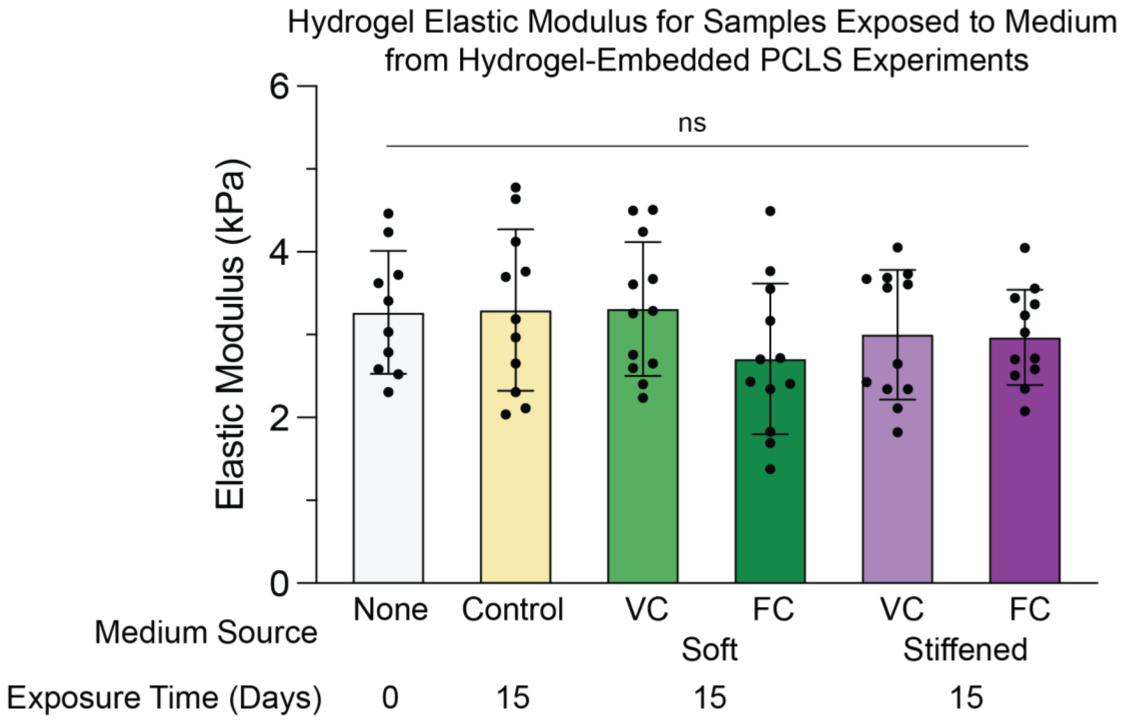


**Supplemental Figure 4.** Supernatant collected from PCLS embedded hydrogel samples suggest that fibrosis cocktail exposure and dynamic stiffening may cause hydrogel degradation and changes in elastic modulus over time. Supernatant was collected from each sample condition and then individually added to acellular soft hydrogels so the elastic modulus could be measured using rheological characterization at discrete timepoints. Columns represent mean ± SEM, n=10-12 with technical replicates. Statistical significance was determined by an ordinary one-way ANOVA test with Holm-Sidak’s multiple comparisons test, ns = not significant.

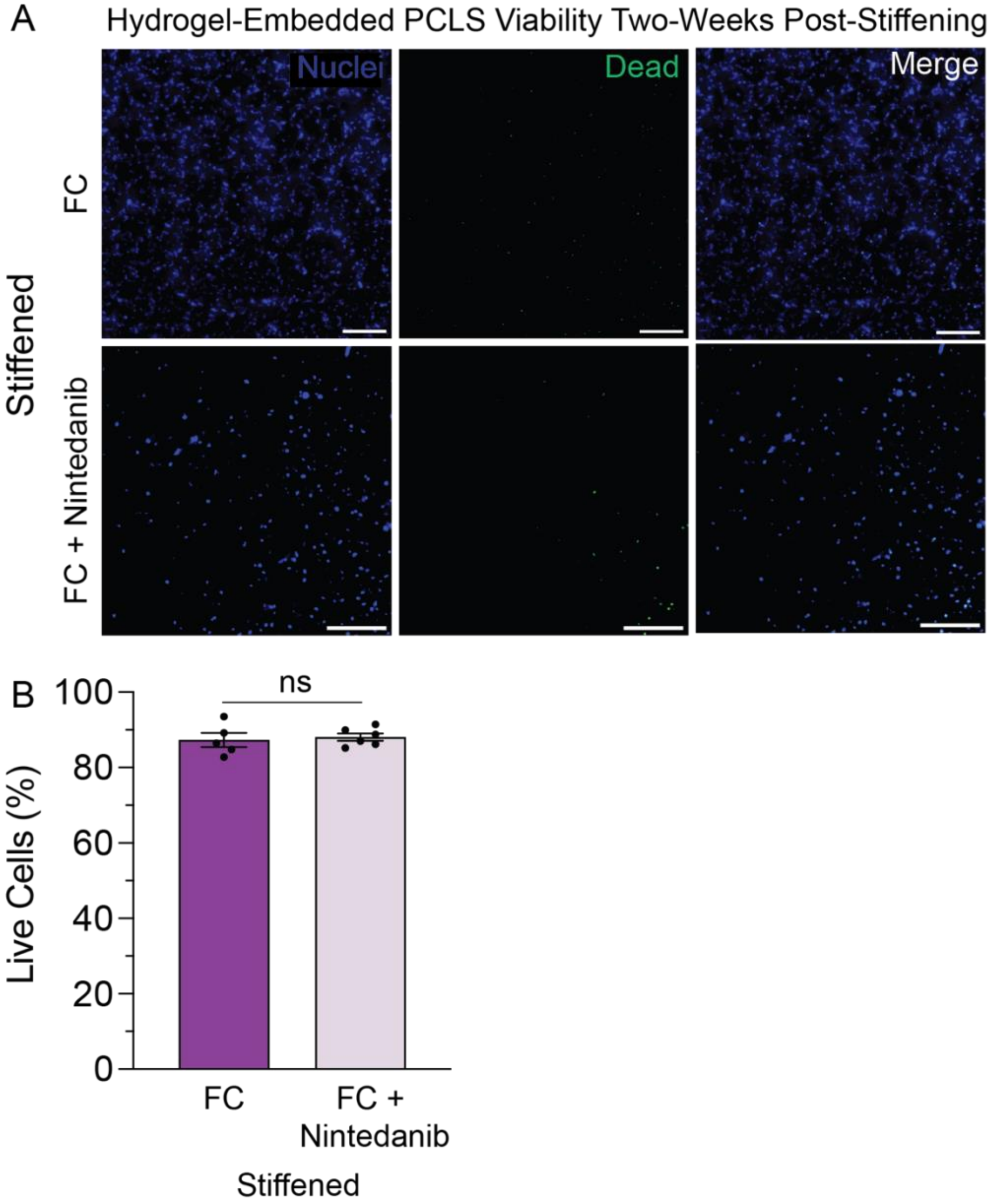


**Supplemental Figure 5.** Quantified live/dead images support that average PCLS cell viability was not negatively impacted by Nintedanib treatment. A) Representative maximum intensity projections of live/dead image z-stacks from stiffened samples. Images show total cell nuclei (blue) and dead (green) cell staining within two weeks post-stiffened hydrogel-embedded PCLS samples that were exposed to either the fibrosis cocktail (FC) or both the FC and Nintedanib. Scale bar, 100 $\mu$m. B) Live/dead quantification results support that the average PCLS cell viability remained above 80% two weeks post-stiffening in stiffened constructs. There was no statistical difference between the samples treated with only the FC and both the FC + Nintedanib. Columns represent mean ± SEM, n=5-6. Symbols represent technical replicates, and statistical significance was determined by an unpaired t-test with Welch comparison test, ns = no significance.

**Supplemental Table 1.** Modified 25% AHA concentration medium formulation per 25 mL.

| Component | Manufacturer/Catalogue Number | Volume ($\mu$L) |
|---|---|---|
| DMEM | Fisher Scientific 21013024 | 24,407.7 |
| 50 mM stock of L-azidohomoalanine (AHA) | Vector Laboratories CCT-1066-25 | 12.5 |
| 50 mg/mL stock of L-methionine | Sigma-Aldrich M5308 | 5.6 |
| 50 mg/mL stock of L-cystine | Sigma-Aldrich C7602 | 24.2 |
| Glutamax | Thermo Scientific 35050061 | 250 |
| 50 mg/mL stock of Ascorbic acid | Sigma-Aldrich A8960 | 25 |
| FBS | Cytiva SH30071.03 | 25 |
| Penicillin-streptomycin | Cytiva SV30010 | 25 |
| Sodium pyruvate | Thermo Scientific 11360070 | 225 |
